# Visual processing and collective motion-related decision-making in desert locusts

**DOI:** 10.1101/2022.09.19.508462

**Authors:** Itay Bleichman, Pratibha Yadav, Amir Ayali

## Abstract

Collectively moving groups of animals rely on the decision-making of locally interacting individuals in order to maintain swarm cohesion. However, the complex and noisy visual environment poses a major challenge to the extraction and processing of relevant information. We addressed this challenge by studying swarming-related decision-making in desert locust last-instar nymphs. Controlled visual stimuli, in the form of random dot kinematograms, were presented to tethered locust nymphs in a trackball setup, while monitoring movement trajectory and walking parameters. In a complementary set of experiments, the neurophysiological basis of the observed behavioral responses was explored. Our results suggest that locusts utilize filtering and discrimination upon encountering multiple stimuli simultaneously. Specifically, we show that locusts are sensitive to differences in speed at the individual conspecific level, and to movement coherence at the group level, and may use these to filter out non-relevant stimuli. The locusts also discriminate and assign different weights to different stimuli, with an observed interactive effect of stimulus size, relative abundance, and motion direction. Our findings provide insights into the cognitive abilities of locusts in the domain of decision-making and visual-based collective motion, and support locusts as a model for investigating sensory-motor integration and motion-related decision-making in the intricate swarm environment.

## Introduction

A fundamental aspect of all instances of collective motion is that of individual repeated decision-making [1–3]. This, in turn, is both driven by and relies on local interactions among the constituent agents, requiring each agent to obtain information about its surrounding social environment [4]. The consequent formation and maintenance of this distinctive form of synchronized movement is understood to be beneficial to the participating individuals [5–7].

A quintessential example of the above process is displayed by the desert locust, *Schistocerca gregaria* (Acrididae). When in the gregarious phase, they collectively move in huge dense marching swarms ([8][9], figure 1A). Locust swarming is commonly accepted as heavily relying on visual perception [10]: each individual locust, with limited visibility amidst an unpredictable terrain, and an intricate, continuously changing social environment, must engage in repeated and dynamic decision-making to avoid getting derailed, while at the same time sustaining the collective motion. This can be translated into a two-layer process: the continuous extraction of the (unknown) state of the social surroundings from the input received by the sensory system (i.e. the eyes); and sensory-motor integration to facilitate the appropriate motor response. Different approaches, ranging from mathematical modeling to studying synchronization in small groups of locusts in laboratory settings, have been employed in the study of swarming behavior in the desert locust [11–14]. However, our understanding of swarm formation and maintenance is still far from complete, partly due to a lack of answers to some fundamental questions regarding decision-making at the individual level.

**Figure 1.**
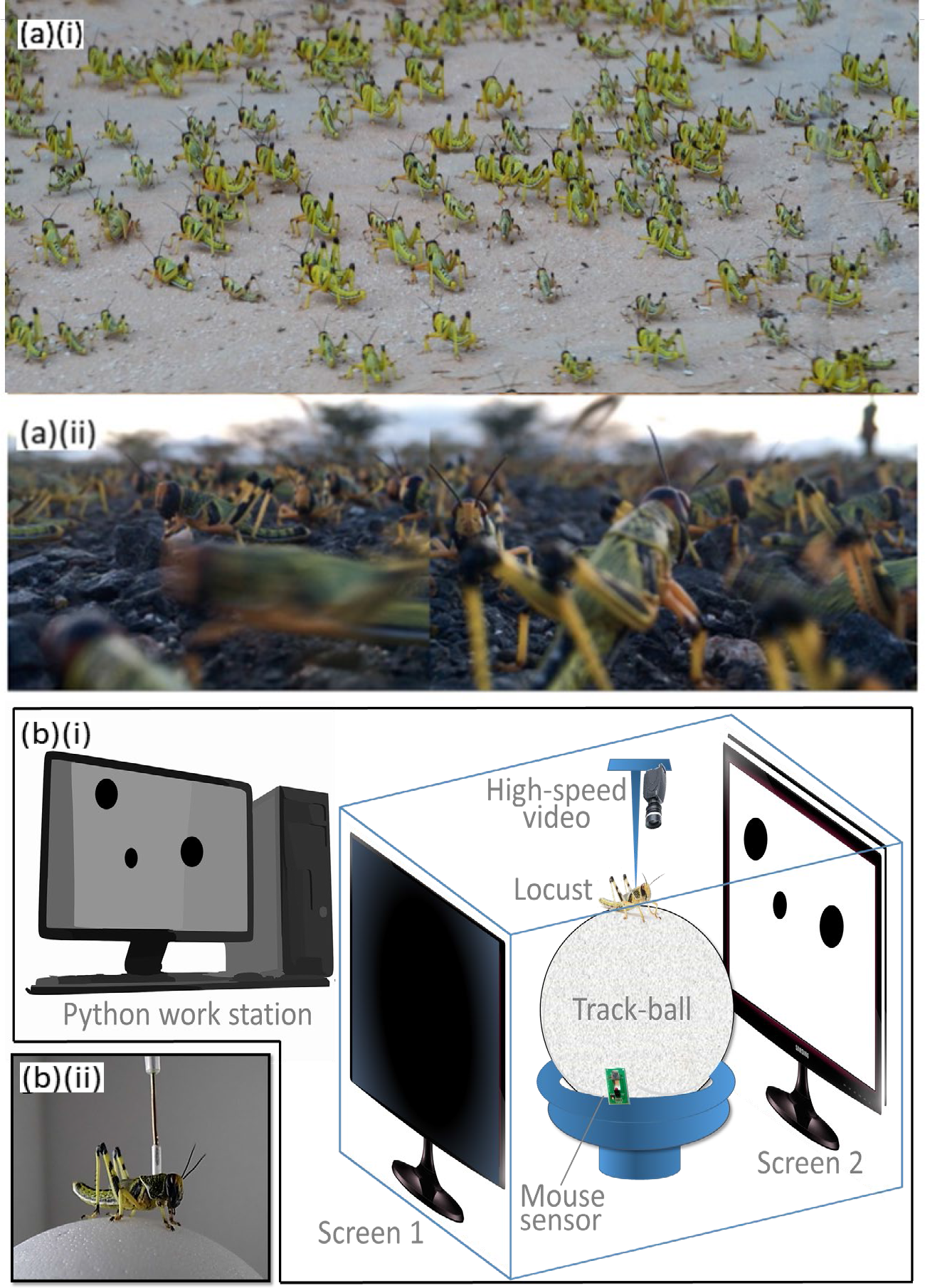
(a) Swarming desert locust nymphs. View from above (a)(i) (Photo taken in Isreal Negev Desert, 2013) and a composit image from the typical height of a locust (a)(ii). (Photos taken in Kenya in 2020 by Inga Petelski). (b)Experimental set-up: complete set-up (b)(i) and individual tethered locust (b)(ii). Individual locust was tethered, with a fixed heading, to an airflow suspended trackball. Random dot kinematograms were presented on two parallel LCD screens. High-speed video camera and a mouse sensor were used for behavioral tracking.

The swift extraction and processing of relevant information from a changing, complex sensory environment presents a critical challenge [15], especially in the noisy and cluttered visual surroundings of a locust swarm. Insects may adopt a range of strategies to increase the efficiency of information perception and processing by reducing the information load [16]. Such strategies include filtering relevant stimuli [17], categorizing the targets [18], and generalizing visual patterns [19]. Filtering relevant visual stimuli, for example by employing a “matched filter” in the visual modality, can reduce the amount of information that needs to be processed [16,20]. In dragonflies and hoverflies, for example, small target detectors are specifically tuned to objects that constitute only a 1–3° angle of the visual field [21,22]. The filtering may occur at different levels of stimuli processing and vary with the ecological relevance of the stimuli. The insect’s nervous system can then channel its resources into performing essential computations, even if complex, in order to extract the task-relevant visual information at low energetic cost [23]. In the case of the desert locust, we hypothesize that, during collective-motion-related visual processing, the locust identifies and extracts relevant stimuli – swarming-related visual cues – from the overall visual scenery, based on a subset of visual features, enabling swift and appropriate decision-making. It is possible that a matched filter for walking speed is used to recognize marching conspecifics; while filtering based on the coherence of the moving group, as inferred from a subset of the swarm, might be used to estimate the overall direction of the swarm.

An additional difficulty imposed on information gathering can arise from the presence of multiple relevant competing inputs [24–26]. In this case, reducing the information load can also be achieved through selective attention – the ability to focus on one type of preferred stimulus while ignoring other perceivable ones [16,27]. Although selective attention remains arguable in the context of insects, the much related key capability to discriminate among different stimuli based on shape, color, and pattern orientation has been observed in honey bees and bumblebees [28–31]; as well as in fruit flies, which show anticipatory behavior consistent with selective attention to the tracked visual stimulus [32,33].

Desert locusts exhibit a characteristic pause-and-go motion, with pause duration correlated with a high probability of turning to change direction [34]. We can thus refer to the locust collective motion as comprising a series of repeated decisions taken by the individuals in the group [13]. Additionally, the decision-making process itself can be considered as a problem of vector selection, including a choice between continued standing or initiating walking, and a choice of direction. Observed variations in the fraction of time spent walking, and particularly in pause duration and the subsequent change in direction, in response to different visual stimuli, can thus offer valuable insights into the locust decision-making process.

We have previously shown that a specific motion-sensitive descending interneuron (one of many behaviorally-relevant descending interneurons (DINs, e.g. [35]), the descending contralateral movement detector (DCMD), conveys information relevant to the locust response to small, slow moving objects (such as other marching locusts [13] and see also [36], [37]). Furthermore, this pathway was shown to demonstrate density-dependent phase-related differences [13], [38], manifested in gregarious locusts being better suited than solitarious ones to the repeated decision-making, and thereby facilitating and coordinating the marching behavior of the swarm. Monitoring the DCMD response to various swarming-related visual stimuli may offer some insights into the neural mechanisms behind the decision-making process under focus in this study.

Here we explored swarming-related decision-making at the behavioral level in *S. gregaria* nymphs, by analyzing different aspects of the individual locust’s walking behavior. These served in our investigation of the role of visual feature recognition and discrimination as possible underlying mechanisms in decision-making. A complementary preliminary electrophysiological study of the processing of visual-motion inputs, relevant to the dynamic interactions between the individuals in a marching swarm, has lent further support to our hypotheses.

## Methods

### Animals

All experiments were carried out using V^th^-instar larvae of *S. gregaria*, taken from our high-density, gregarious phase locust lab-colony at the School of Zoology, Tel Aviv University (rearing conditions were as recently described in [12].

### The experimental setup

Individual locusts were tethered in a fixed (forward) head direction, via a 1 cm long clear vinyl tube attached to their pronotum with epoxy resin, in a natural-like typical walking posture, above an airflow-suspended Styrofoam trackball, illuminated from above with LED lights. The ball was decorated with an irregular black over white pattern in order to facilitate the tracking of its movement. Two parallel LCD screens, 30 cm apart, were positioned one on either side of the locust, allowing the presentation of controlled visual stimuli, while carefully monitoring the locust’s behavioral responses and movements of the ball by a high-speed video camera (figure 1B). Experiments started after one hour of acclimation of the locust to the tether. In two sets of behavioral experiments the locusts’ responses were monitored using FicTrac [39], a computer-vision tracking software that determines the angular position of the ball for each frame. In an additional set of experiments, an optical mouse sensor was further utilized to record the movement of the ball. The behavioral setup was complemented by a corresponding electrophysiological setup, enabling the recording of the neural responses of the locust DCMD interneurons to (similar) controlled visual stimuli (see Electrophysiology section below).

### The visual stimuli

Visual stimuli, designed using the Python programming language and PsychoPy (an open source software package; [40]), were presented in the form of random dot kinematograms (RDK) of black dots on a white background, at a maximum contrast of 100%. We chose RDK following previous reports of utilizing such stimuli for testing multiple target processing, and specifically motion perception [41]. Unless stated otherwise, the RDK comprised 40, 1.2 cm diameter dots, corresponding to a subtended visual angle of 6.86° on the insect’s eye (within the known size of the locusts). Each visual stimulus was presented for 60 seconds.

We first presented the control stimuli: (1) blank (white screen), and (2) still dots on a white screen. Next, we conducted a set of different behavioral experiments to investigate the tethered locust’s response to the following different tentative features of swarming-related visual stimuli:

#### Direction of motion

The RDK comprised fully coherent, 5 cm/s moving dots, simulating a coherently moving locust swarm. Three types of stimuli were used, each with a different direction of motion: (1) both screens showing dots aligned with the direction of the tethered locust’s heading; (2) both screens showing dots in a direction 180° to the tethered locust’s heading; and (3) one screen showing aligned dots and the other with dots moving in the opposite direction.

#### Motion speed

Tethered locusts were presented with dots moving with 100% coherence on both screens, aligned with the tethered locust’s heading, and at graded speeds. The tested motion speeds were 1, 3, 5, 10 and 15 cm/s, which cover a marching locust’s speed range, as measured previously [12].

#### Coherence level

Tethered locusts were presented with dots moving on both screens at a motion speed of 5 cm/s and graded coherence levels: a fraction of the dots moved in alignment with the locust’s heading while the remaining dots each moved in a random direction. Coherence levels tested were 0 (all dots moving in different random directions), 0.1, 0.25, 0.5, and 1 (all dots aligned with the locust’s heading direction).

#### Competing stimuli

A fourth experiment was conducted to investigate situations of competing stimuli, i.e., decision-making in the presence of conflict. First, a more quantitative type of conflict was presented to the locusts: 2/3 of the dots on each screen moved in one direction, either aligned with or opposite to the tethered locust’s heading, while the remaining 1/3 moved in the other direction. Next, a size difference was added (size mimicking proximity differences): the 1/3 dots moving in the opposite direction to the 2/3 were also double the size of the latter (2.4 cm diameter).

### Behavioral analysis

The rotation angle, the difference between two angular positions of the trackball in subsequent frames, was used to analyze the locust motion parameters. A motion threshold was determined based on the extent of the rotation angle. A locust was considered to be moving if the threshold was crossed for at least 10 consecutive frames. Pausing was determined if the same threshold was not crossed for at least 20 consecutive frames. Based on these indices, we calculated the fraction of time spent walking (walking fraction) and the average pause duration. A sideways motion (positive and negative) threshold was determined based on the direction of the trackball rotation. A locust was considered to be moving sideways if this threshold was crossed for at least 10 consecutive frames. The coordinate positions from the optical mouse sensor were used to determine the walking parameters by calculating the walking distance, using the Cartesian formula.

### Electrophysiology

Dissection and electrophysiological procedures followed Ariel et al., (2014). Briefly, following CO_2_ induced anesthesia, the legs of the nymphs were removed and a silver hook electrode was positioned around the ventral neck connectives for extracellular recording of DCMD activity. The locusts were positioned above a plastic platform in the same position and posture as on the airflow-trackball. The experiments were performed using RDKs with graded speeds and coherence levels similar to the behavioral experiments, and also using graded sizes of 0.4, 0.8, 1.2, 1.4, 2.2, 5.5 and 6.8 cm diameter. Each stimulus was presented for 20 seconds, with an inter-trial period of 1 min. The DCMD action potential times, number, and frequency were analyzed.

## Results

### The locusts respond to swarming-related visual stimuli with a preference to maintain their heading

Locusts tethered in our setup exhibited the typical pause-and-go motion pattern even in the absence of moving visual stimuli (see also (10,34)). Similar walking kinematics (manifested in pause duration and walking fraction) were measured in response to the white background only and to the background with motionless dots (Fig. S1). When testing the effect of moving stimuli, the response of the locusts comprised several clear time-dependent features (Fig. S2). Specifically, when the direction of the stimulus motion on one or both screens was opposite to the tethered locust’s head direction, the demonstrated behavioral response was not consistent throughout the stimuli, but became exhausted prior to termination of the stimuli. This time-limited response was probably due to the open-loop nature of our experiments, i.e., to the fact that the locust’s response did not induce any (expected) directional change in the incoming visual inputs. Consequently, we limited our comparative analysis of the different visual-stimuli-induced behavioral responses to the first 40 seconds of each trial only. During this consistent and robust “responsive time-interval”, the locusts attempted to align themselves with the direction of motion of the dots, and/or to join the motion (figure 2).

**Figure 2.**
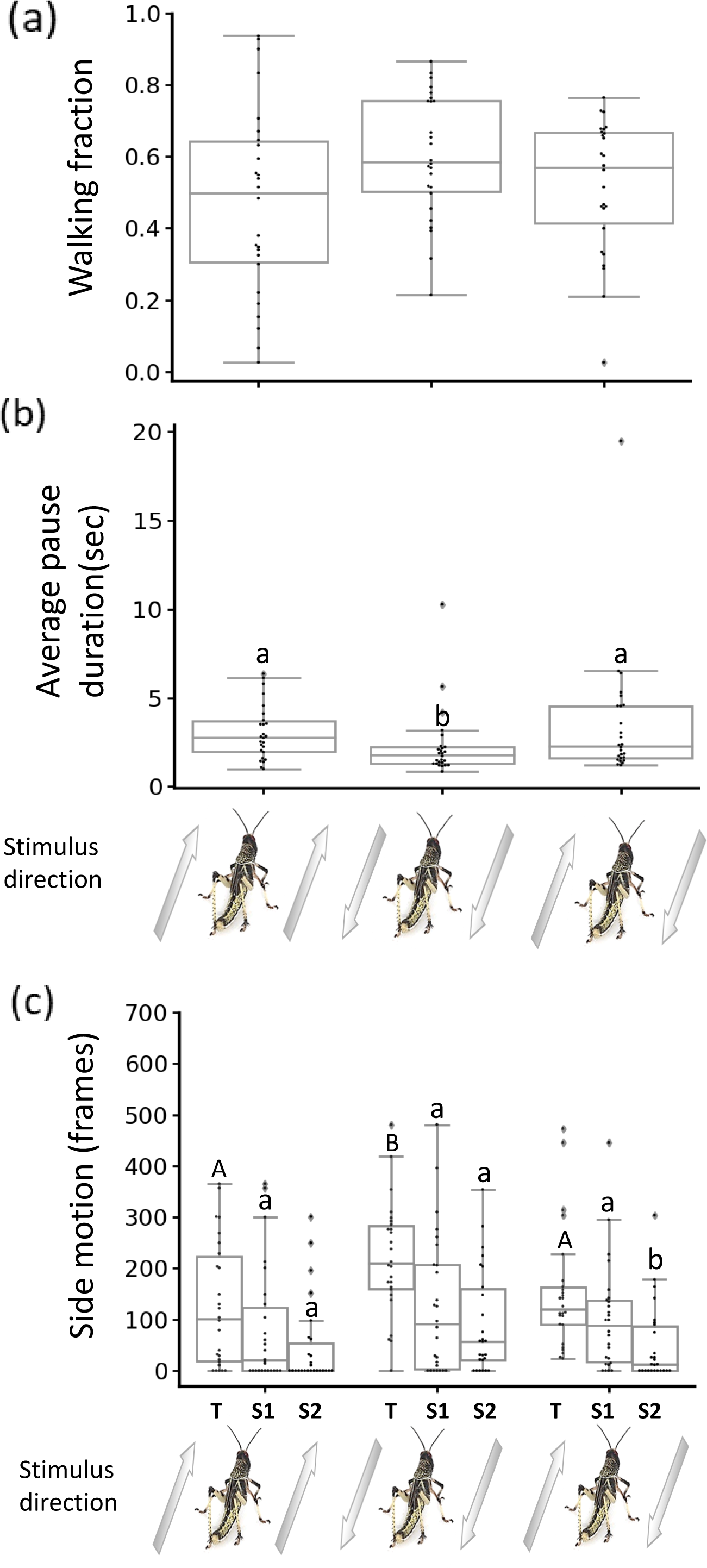
Locust response to swarming-related visual stimuli. Walking fraction (a), average pause duration (b) and side motion (c) in response to different motion directions of the stimuli. Arrowheads represent direction of motion relative to locust heading. Each point represents data from a single locust (n = 26) Gray lines denote the median. Boxes show the interquartile range (25th to 75th percentiles). Whiskers include points up to 1.5 times the interquartile range. (a)-no significant difference between different stimuli (b)-Different letters represent statistical differences. (c)- S1-side motion versus one side, S2-side motion versus the other side and T=total side motion (S1+S2). Different capital letters represent statistical differences in total side motion between different types of stimuli; different lower case letters represent statistical differences between different sides (S1 and S2) of the same stimulus.

When characterizing the behavioral response to stimuli moving on both screens in the locust’s heading, compared to both screens showing stimuli moving in the opposite direction, the latter generated a significantly decreased average pause duration (figure 2b, n=26, Friedman test, p<0.05, Dunn’s multiple comparisons test, p<0.05) and significantly increased overall side motion (figure 2c, n=26, Friedman test, p<0.001, Dunn’s multiple comparisons test, p<0.01). When each screen displayed a different direction of motion, one aligned with the locust’s head direction and the other opposite to it, no significant difference was observed in the above-noted parameters between this condition and the two others. However, the locust’s side motion towards the monitor displaying moving dots in a direction aligned with its heading was significantly higher compared to its side motion towards the other monitor (figure 2c, n=26, Wilcoxon matched-pairs signed rank test, p<0.05). No such preference for motion towards a specific side was noted when both screens displayed stimuli with the same direction of motion. Overall, these findings confirm the swarming-related nature of our controlled stimuli, i.e., the locusts clearly attempted to swarm alongside or to join the controlled visual stimuli presented in our experimental setup, demonstrating a preference towards stimuli that were aligned with their initial heading.

### Clear thresholds are demonstrated in the response to swarming-related visual stimuli at both the individual conspecific and the group level

The next feature investigated for a possible effect on the locust’s behavior was motion speed. As can be seen in figure 3a, a clear dependence and a clear speed threshold were demonstrated: in response to stimuli with motion speed greater than 3 cm/sec, significantly higher walking fractions (figure 3a(i), n=15, Kruskal-Wallis test, p<0.0001) and shorter pause durations figure 3a(ii), n=15, Kruskal-Wallis test, p<0.0001) were observed, compared to the response to dots moving at speeds below this threshold.

**Figure 3.**
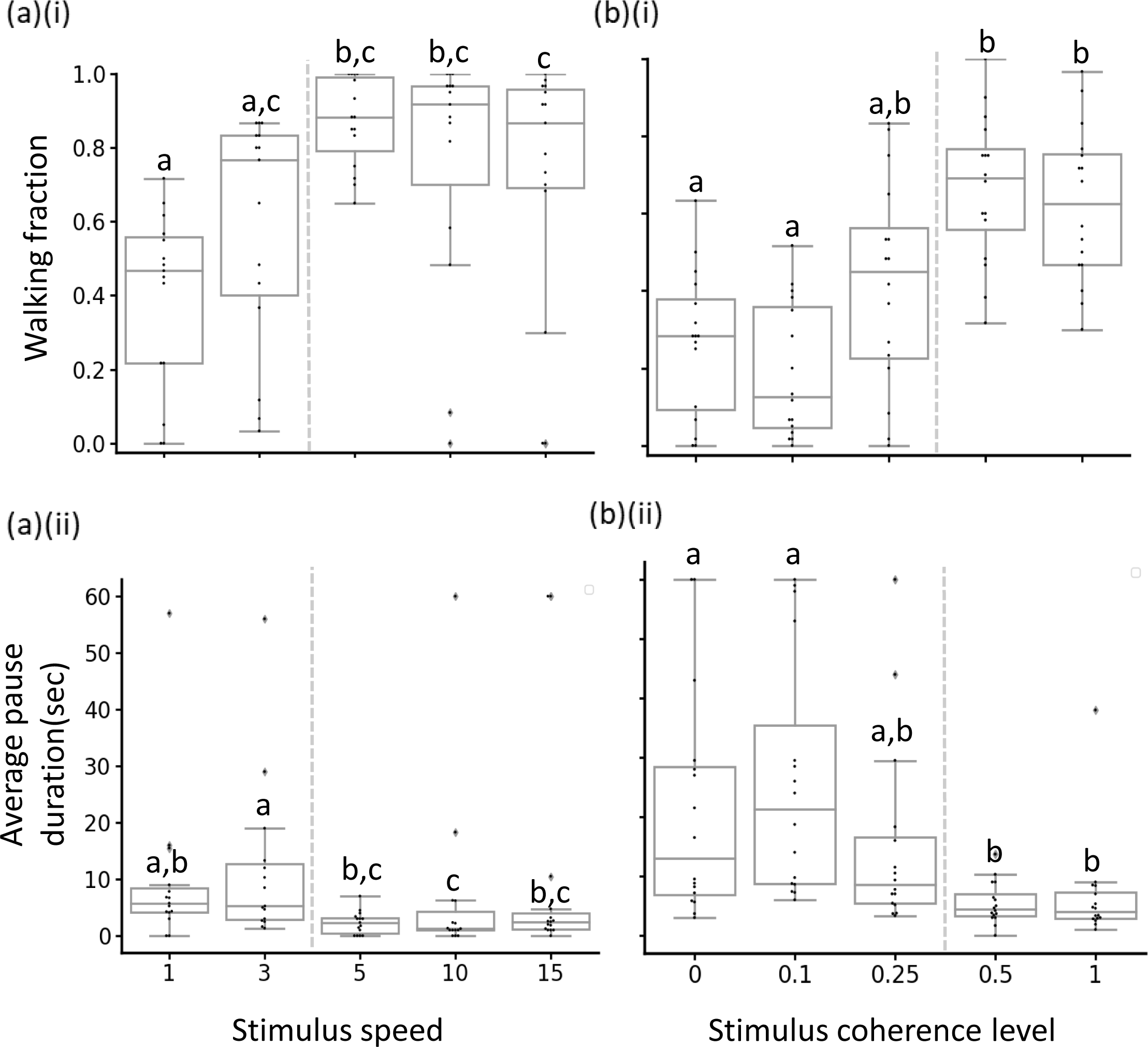
Behavioral thresholds in response to swarming-related visual stimuli. Average pause duration (a)(i) and walking fraction (a)(ii) in response to different stimulus motion speeds (n = 15). average pause (b)(i) and (b)(ii) in response to different stimulus coherence levels (n = 16). Each point represents data from a single locust. Gray lines denote the median. Boxes show the interquartile range (25th to 75th percentiles). Whiskers include points up to 1.5 times the interquartile range. Points that are more than 1.5 times the interquartile range away from the bottom or top of the box are outliers. Different letters represent statistical differences. Gray dashed line indicates location of behavioral threshold.

Maintaining the speed of all the moving dots above the demonstrated threshold, and changing the coherence level among the presented dots, revealed a second decision rule based on yet another threshold (figure 3b): in response to stimuli with coherence level above 25%, the locusts exhibited significantly larger walking fractions (figure 3b(i), n=16, Kruskal-Wallis test, p<0.0001) and significantly shorter pause durations (figure 3b(ii), n=16, Kruskal-Wallis test, p<0.0001) compared to their response to stimuli with coherence levels below this threshold. It is important to note that while both the speed and the coherence level are characteristic features of the visual inputs in a marching swarm, the former is a feature of each individual group member, contributing to the collective motion of the swarm; while the latter is a characteristic of the collective, or a group-level trait, reflecting the common direction of motion.

### The locust response to complex moving visual stimuli

As noted, the visual environment within a locust swarm is an intricate and noisy one, intriguing us to investigate locust response to complex and conflicting stimuli (figure 4). First, locusts were presented with visual stimuli comprising two groups: the 2/3 group of dots moved in one direction and the 1/3 group of dots moved in the opposite direction All dots moved at similar speeds, above the speed threshold demonstrated previously. Next, a size difference was introduced, such as the dots in the minority group being twice the size of those in the majority group. When 2/3 of the dots were moving in a direction aligned with the locust’s heading and the smaller group of dots were moving in the opposite direction, no significant effect was observed in the locust’s response following an increase in size of the dots in the smaller group. However, when the majority of dots were moving in the opposite direction to that of the locust’s heading, changes in locust kinematics were noted. When all the dots were equal in size, the locust’s walking fraction significantly decreased (figure 4a, n=19, Friedman test, p<0.05, Dunn’s multiple comparisons test, p<0.05). This was possibly due to the conflict between relative abundance (2/3 of dots moving in the opposite direction) and the preferred motion direction still present in the remaining 1/3. Doubling the size of the dots in the 1/3 group, moving in the direction of the locust’s heading partially restored walking fraction and significantly increased pause duration (figure 4b, n=19, Friedman test, p<0.01, Dunn’s multiple comparisons test, p<0.05). This specific complex visual stimulus required more intensive information processing by the locust, demonstrated by the larger pauses.

**Figure 4.**
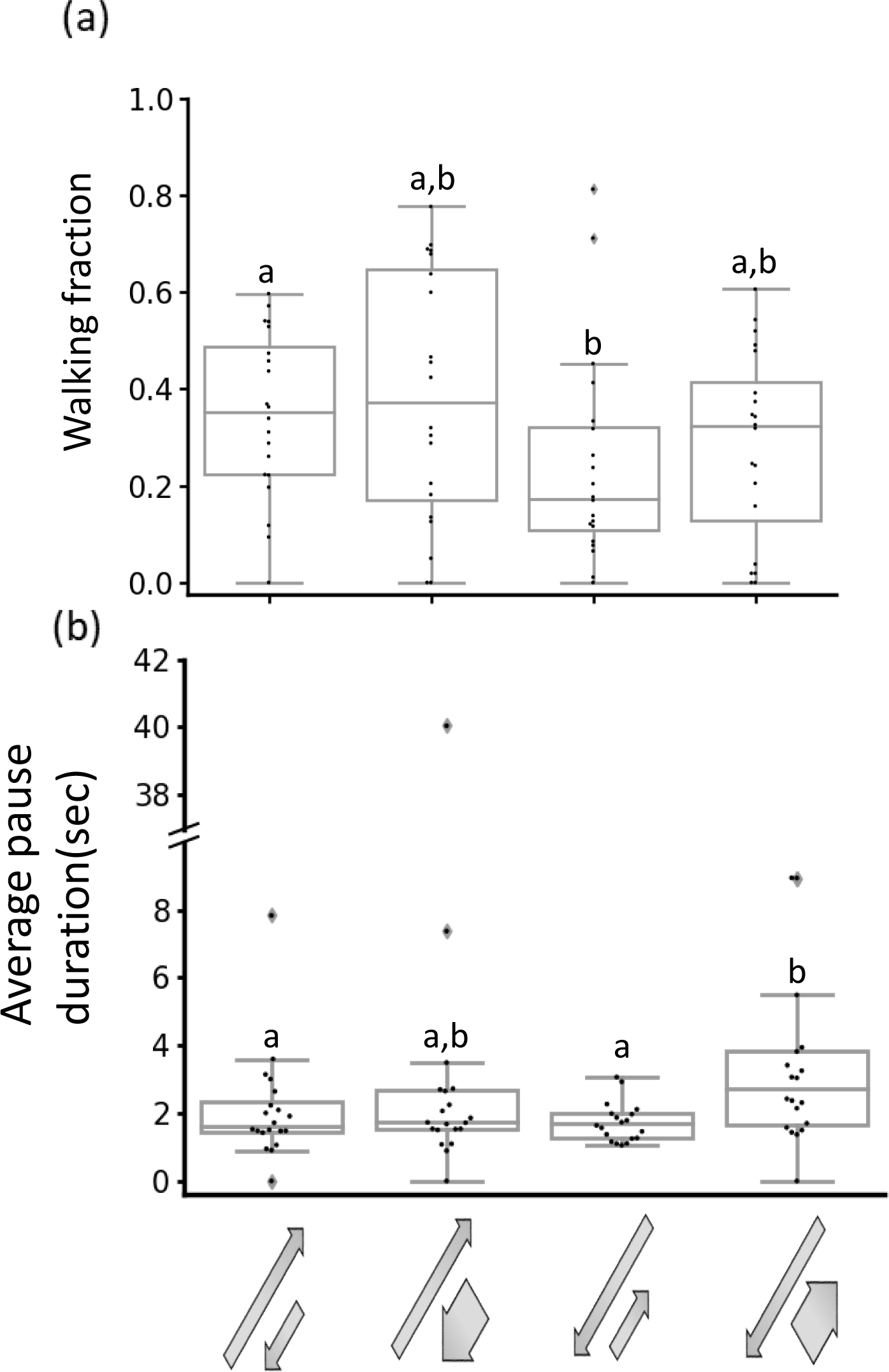
Size-direction-abundance interplay in response to complex visual stimuli. Walking fraction (a) and average pause duration (b). Arrowheads represent direction of motion. Arrow length represents relative abundance (long arrows = 2/3 of dots, short arrows = 1/3 of dots). Arrow width represents dot size (wide arrows – larger dots). Each point represents data from a single locust (n = 19) Gray lines denote the median. Boxes show the interquartile range (25th to 75th percentiles). Whiskers include points up to 1.5 times the interquartile range. Different letters represent statistical differences.

Overall, these findings reveal intricate interactions between stimulus number, size, and direction, which together affect the locust decision-making process.

### Neurophysiological correlates to the responses to swarming-related visual cues

Further exploration of the sensory-motor processing of swarming-related visual cues was conducted through a series of neurophysiological experiments.

Based on the behavioral observations, we expected our different visual stimuli to induce variable neuronal responses, depending on the stimuli motion speed, coherence level, and dot size. This hypothesis was tested by studying the response of the DCMD interneuron, a key participant in a well-described motion sensitive visual pathway [42,43], to similar types of stimuli as above. The DCMD has been mostly studied in the context of looming stimuli. Hence, it should be noted that the responses observed and monitored in our experiments differ from those of the typical looming response (figure 5). The DCMD firing rate in response to control stimuli (white background and still dots) was similar to its spontaneous firing rate reported in previous studies [13]. Manipulating the characteristics of swarming-related visual stimuli thus induced different responses:

**Figure 5.**
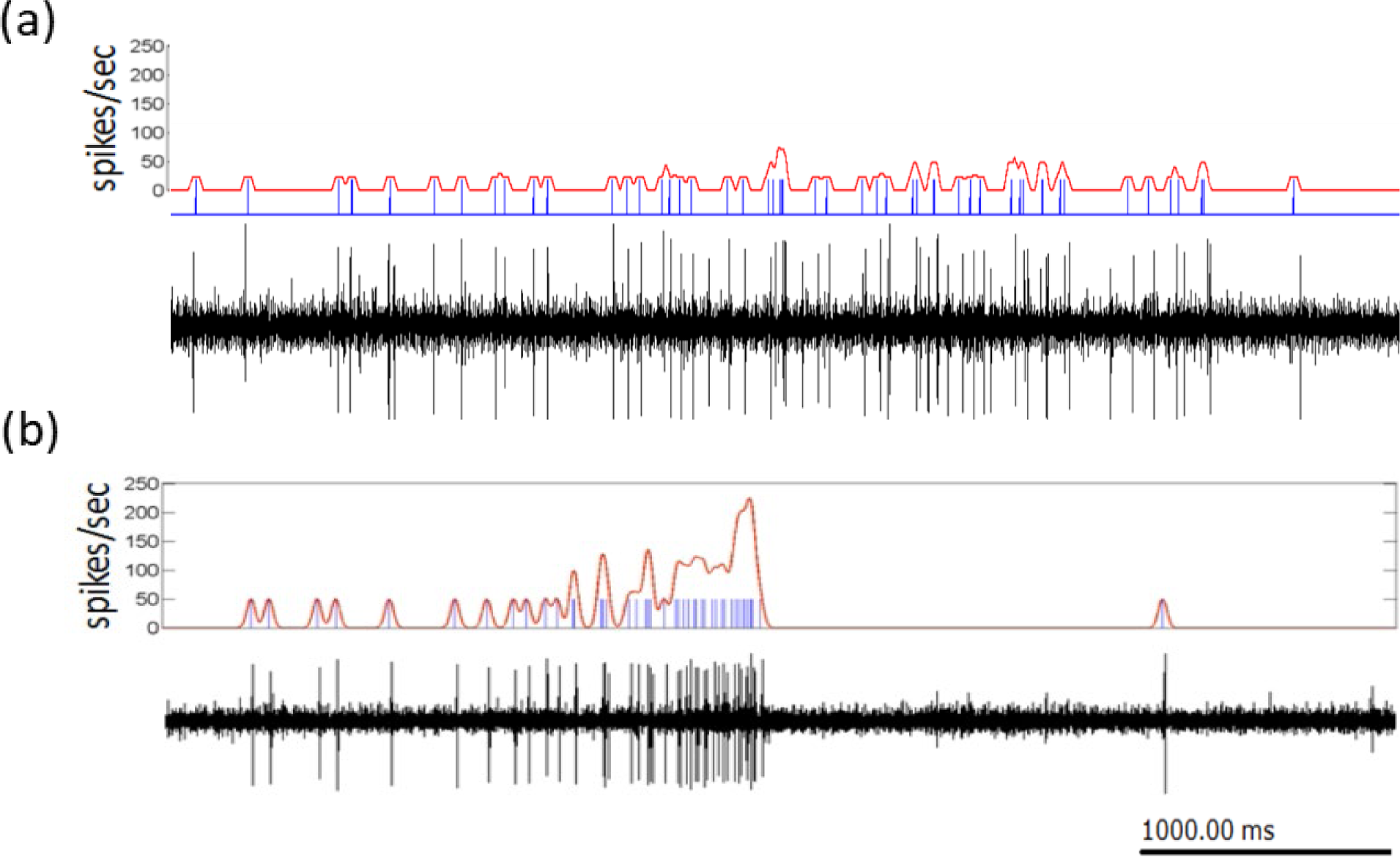
Typical response of the DCMD to swarming-related (a) and looming (b) stimuli. DCMD spike occurrence times (blue) were extracted from the extracellular recordings (black). Individual raster trials were then smoothed with a 20 ms Gaussian window and an evaluation of the instantaneous firing rate (red) was calculated. Recording (a) is the responses to swarming-related visual stimuli with motion speed of 5 cm/s and recording (b) is a response to looming stimulus (modified from Ariel et al. 2014).

#### Motion speed

We found a clear dependence of the DCMD firing rate on the speed of the moving dots. A significant difference was seen between the slowest tested motion speed – 1 cm/s, and the fastest one – 15 cm/s (figure 6a, n=7, Friedman test, p<0.001, Dunn’s multiple comparisons test, p<0.001), comprising two extremes within the speed range of marching locusts [12]. The DCMD’s responses to visual stimuli moving at intermediate speeds did not significantly differ from each other.

**Figure 6.**
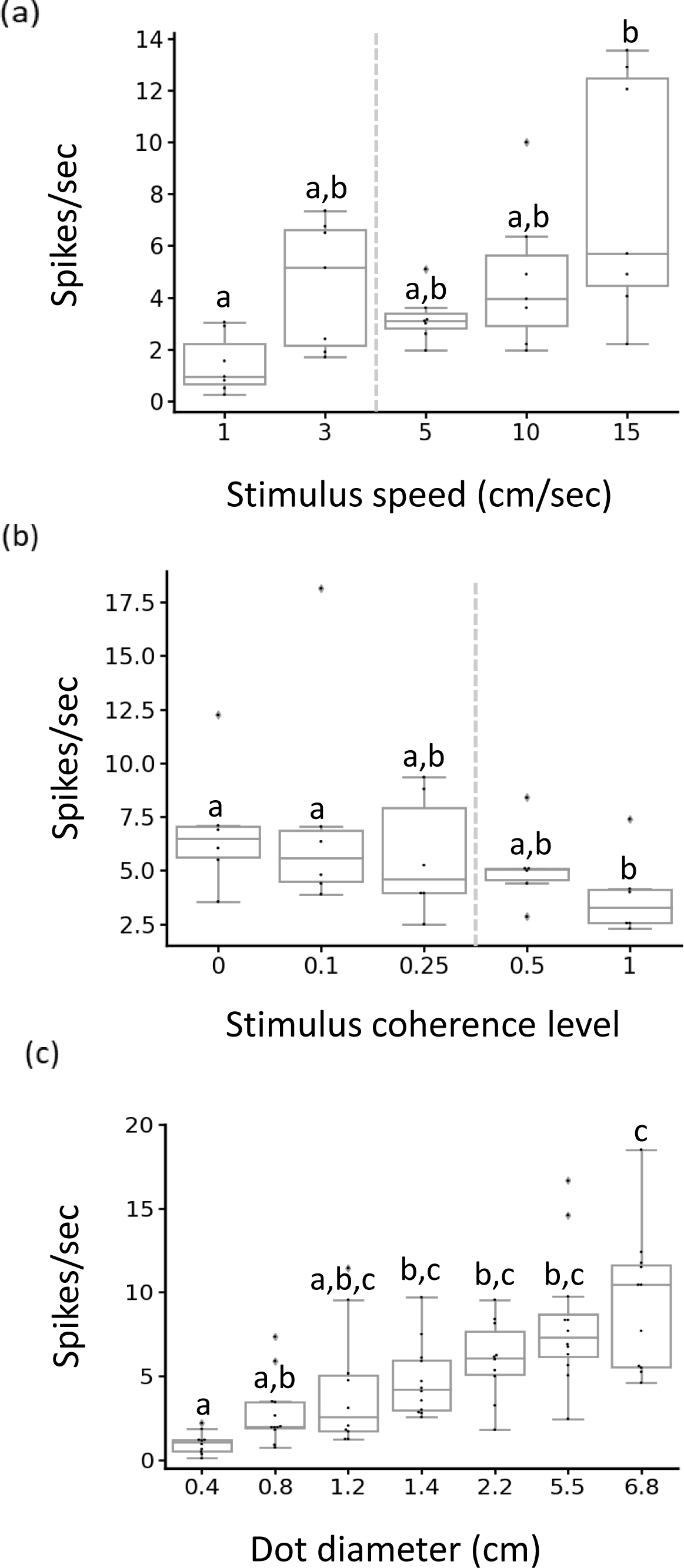
DCMD response to different motion speeds (a), coherence levels (b) and dot size (c) of swarming-related visual stimuli. Each point represents data from a single locust((a) n=7, (b) n=6, and (c) n=10). Gray lines denote the median. Boxes show the interquartile range (25th to 75th percentiles). Whiskers include points up to 1.5 times the interquartile range. Different letters represent statistical differences. (a) and (b) - Gray dashed line indicates value of behavioral threshold.

#### Coherence level

Low coherence levels elicited high DCMD firing rates, with the DCMD response declining with the increase in motion coherence level from 0/1 to 1 (figure 6b). DCMD firing rate with coherence levels of 0% or 10% was significantly higher compared to that in response to 100% coherent stimuli (figure 6b, n=6, Friedman test, p<0.01, Dunn’s multiple comparisons test, p<0.01).

#### Size effect

Testing the DCMD response to swarming-related moving stimuli comprising dots of different sizes, revealed a size-dependent firing rate: a significant difference was noted between dots with a diameter of 0.4 cm and those with a diameter of 2.2, 5.5 or 6.8 cm (figure 6c, n=10, Kruskal-Wallis test, p<0.0001, Dunn’s multiple comparisons test, p<0.051). A graded increase in firing rate was seen with the increase in size.

Overall, our neurophysiologic investigation revealed that the DCMD was sensitive to different speeds, coherence levels, and sizes of swarming-related visual stimuli, in a similar though not identical manner to that revealed in our behavioral experiments.

## Discussion

Sensory information has a crucial role in ecological decision-making [15]. In order to enable sensory processing to be swift and context-appropriate, organisms are required to identify and extract highly specific, behaviorally relevant, signals from their surroundings [44]. Different strategies for rapidly coping with a visually cluttered environment have been suggested in previous studies of different organisms engaged in vision-based collective motion [13,45–47]. Flocking birds were reported to consider visual information from a fixed number of influential neighbours (i.e., a topological range; [48]). Zebrafish rely on visual occupancy for direction choice [45], and use bout-like movements for conspecific recognition [49]. Collectively moving *Drosophila* larvae depend on the number of conspecifics and cues related to their unique visual kinematic for decision-making [50]. Beyond the principal role of motion sensitivity in maintaining synchrony during collective marching [11,12], only very limited knowledge is available regarding how locusts utilize visual-sensory cues for swarming-related decision-making amidst their highly challenging visual surroundings.

Our findings have identified specific characteristics of the behaviorally-relevant visual inputs affecting decision-making in desert locust nymphs. Moreover, we show, to the best of our knowledge for the first time, that locusts can extract collective-motion relevant information at both the individual conspecific level (i.e. speed) and the group level (coherence or common direction), possibly by means of filtering and discrimination. While filtering can reduce the information processing load at the very first stage by differentiating relevant from non-relevant stimuli and ignoring the latter, discrimination can aid the extraction of information from the relevant stimuli and subsequently facilitate critical decision-making.

Desert locust nymphs walk at an average speed of ∼5 cm/sec [12]. In response to coherent stimuli moving at non-zero speeds below 3 cm/sec, the tethered individuals exhibited longer pause durations and shorter walking fractions, possibly reflecting longer decision-making time due to a mismatch between dot speed and the expected behaviorally relevant conspecifics’ speed. Hence, the locusts employed filtering at the level of the characteristic of the individual. In natural settings, such a clear speed threshold may be exploited to recognize marching conspecifics, such that anything moving at a speed below the threshold is ignored. This behavioral threshold was discovered in the V^th^-larval instar nymphs (corresponding to walking speed at this stage). Since locust collective marching appears early on and is maintained throughout the different developmental (larval) stages [51], and as development is accompanied by a marked change in size as well as in walking speed, an interesting point for future research would be that of developmental plasticity within the observed speed threshold.

The relatively limited walking behavior demonstrated in response to dots moving in a non-coherent fashion (a non-decisive state manifested by unusually long pause durations), could have resulted from a lack of appropriate relevant information at the level of the group movement pattern. Importantly, our findings indicate that the locusts perceive a complete absence of motion cues (i.e., still dots) differently to that of an absence of conclusive information in the motion cues (i.e., non-coherently moving dots). This could be reflected in gregarious locusts continuing to walk even when alone (albeit with altered kinematics [14], but tending to pause and wait when surrounded by conspecifics moving randomly, e.g., early morning at the initial organizing stages of a swarm in a natural setting[51,52]. This ability to extract the trajectory of the surrounding locusts seems to be a fundamental characteristic of gregarious-phase locusts, instrumental for the decision-making in enabling the collective motion of locust swarms.

In his review, Warrant [53] discusses different visual matched filters and their important role in the ecology of vision in insects. These include peripheral matched filters for sex (mate), for prey detection and pursuit, and for the physical environment (aspects of the physical terrain). Central visual matched filters include filters for the insect’s own locomotion speed (“fast” or “slow” eyes) and for navigation (the celestial pattern of polarized light). Our above findings suggest yet another possible matched filter that is crucial for locust swarming behavior: this can be referred to as a “social environment” matched filter, or maybe even filters, as we have demonstrated filtering at both the level of the motion of individual neighboring conspecifics as well as of the surrounding group.

Relevant conflicting stimuli increase the difficulty imposed on information processing and decision-making. A conflict can derive from a contradiction in one feature (e.g. opposing motion directions), requiring a single-attribute decision, or from the complex interactions between several features (e.g. direction, abundance, and size), becoming a multi-attribute choice problem [54]. Locusts presented with directionally contradicting but otherwise identical swarming-related visual stimuli, demonstrated a preference for stimuli with motion direction aligned with their own heading. This preference to join in the marching aligned to the locust’s current heading is consistent with the observation that marching locusts in experimental ring-shaped arenas only rarely change direction [11,13]. When different parameters provide conflicting or inconsistent information, it becomes beneficial to assign a higher weight to one over the other. Ants, for example, rank several attributes when faced with a multi-feature problem [55], and the ranking is dynamic in relation to their current situation [56]. When presented with mixed-type stimuli, size affected the locust’s behavior only when in a specific relation to the direction of motion and relative abundance. The interplay between these three parameters significantly affected pause duration, which is the kinematic phase assumed to be dedicated to information processing and decision-making [13,34]. Size may act as a proxy for visual target distance, with nearest neighbors being larger. More attention dedicated to larger dots, although less abundant, indicates that immediate neighbors may influence the decision of an individual more strongly than more distant members of the swarm. This is also consistent with previous reports suggesting a limited functional radius of attention around an individual in a group [13];[14].

The DCMD is only one out of several currently known descending interneurons that take part in motion-sensitive pathways [57,58]. It has been widely researched for its characteristic response to looming visual stimuli, and its function in predator and collision avoidance maneuvers [59–62]. Nevertheless, the motion-sensitive pathway in which it takes part is capable of responding to different, complex types of visual-motion stimuli [37,62]. The DCMD was also shown to demonstrate activity changes with developmental stages [36]. Ariel et al. [13] demonstrated the its ability to convey information regarding motion types similar to those of marching locusts, and a specific “tuning” of the response habituation rate in swarming, gregarious-phase locusts compared to the non-swarming solitarious-phase ones. Hence, the DCMD response to swarming-related visual stimuli, to which locusts demonstrated behavioral responses, can provide information regarding the neurophysiological mechanisms underlying these behavioral responses, and the related decision-making process. The DCMD in our experimental setting demonstrated consistent responses to non-looming visual stimuli, with a clear effect of changes in motion speed, coherence level, and stimulus size. The sensitivity to changes in these specific visual stimulus features emphasizes their importance and supports the involvement of these during visual information processing in swarming-related decision-making.

In its simplest form, decision-making can be understood as the process of selecting between two alternatives. The flexibility of decisions is accepted as a trademark of higher cognition in organisms [63]. Given that cognitive performances and integral abilities are often assumed to be positively correlated with brain size across species, it is no surprise that the miniature brains of insects are believed to limit their computational power and cognitive abilities [64–67]. However, with mounting evidence in support of highly sophisticated behaviors in insects [68], this assumption currently holds little ground. Here, we have provided additional insights into insect cognition by investigating decision-making in a collectively marching insect species. We demonstrate that desert locusts use discrimination, mediated by selective attention, to extract relevant information from a complex, noisy, visual environment. The rules of decision-making (decision rules) in gregarious desert locusts seem to be a function of multiple interacting factors, with the ultimate goal of staying in sync with conspecifics. A differential weightage to parameters such as direction, number, and size of the stimuli was also observed, with conflicting information increasing the difficulty imposed on the decision-making process.

To conclude, locusts utilize different mechanisms that enable them to meet the challenges presented by the overloaded and cluttered visual environment, and that support the sensory perception and integration required for collective motion-related decision-making. These mechanisms constitute an instrumental aspect of their ability to synchronize with conspecifics and maintain the cohesion of the swarm, and thus probably also exist in all animals demonstrating visual-based collective motion. Much further work is required, however, in order to uncover and describe the details of the sensory-motor integration (e.g. the role of feedback from the motor system), to fully elucidate the underlying neurophysiological mechanisms (e.g. additional key neuronal pathways), and to provide insights into related brain-level phenomena, such as the representation of conspecifics and their behavior (speed, direction, etc.), the actual depiction or abstraction of the swarm as a whole, and more.

## AUTHOR CONTRIBUTION

AA conceived the study; IB and PY carried out experiments and data analysis. AA and IB and PY wrote the manuscript

The authors have no competing interests related to this study.

Supplementary material is available online on DRYAD: https://doi.org/10.5061/dryad.jdfn2z3dw

